# Differential collagen crosslinking and network organization creates distinct tissue remodeling patterns in fibrosis and COPD

**DOI:** 10.64898/2026.05.13.724372

**Authors:** M. M. Joglekar, M. Nizamoglu, M. C. Morrison, R. Hanemaaijer, T. Koster, K. Sjollema, T. Borghuis, M. C. Zwager, I. H. Heijink, S. D. Pouwels, B. N. Melgert, N. Gavara, J. K. Burgess

## Abstract

Collagens are key components of the extracellular matrix (ECM) that play a crucial role in maintaining structure, strength, and function of the lungs. Fibrillar collagens are crosslinked by enzymes such as lysyl oxidases and transglutaminases and organized into networks by proteoglycans and glycoproteins. Collagens are the main load-bearing components and along with elastin may impart a non-linear strain hardening behavior to the lung. In disease, collagen crosslinking and organization can be disrupted, possibly due to abnormal levels of enzymes or ECM components. Few studies have examined collagen crosslinking and organization in healthy and diseased human lungs. In this study, alterations in collagen crosslinking and organization were investigated in human lung control, fibrotic and chronic obstructive pulmonary disease (COPD) tissue sections. Ultra-performance liquid chromatography and second harmonic generation microscopy measured pyridinoline crosslinks and the distribution of mature and immature collagens within the decellularized scaffolds, respectively. Fibrotic scaffolds had higher total collagen but less crosslinking per mole of collagen compared with COPD donors. Image analysis by second harmonic generation microscopy showed mature collagens populated airway or blood vessel walls in all three groups and in the parenchyma of fibrotic scaffolds. Immature collagens, on the other hand, were mainly localized to parenchymal regions in control and COPD scaffolds, with fewer immature collagens in fibrotic parenchyma. Additionally, quantification of the mature to immature collagen ratio in defined regions of control and diseased scaffolds showed increased organized collagen in fibrotic tissue. Our study shows that collagen crosslinking and organization are disrupted in fibrotic and COPD lungs and these changes may be compartment specific and can contribute to aberrant mechanical properties of diseased lungs. Our findings highlight that along with total collagen content, collagen crosslinking and organization are equally important while investigating collagen-mediated pathological changes in lung tissue. These changes may have implications for developing ECM-based therapeutics for patients with lung diseases.

## Introduction

Lung structure and function are strongly associated with each other. The intricate architecture of alveoli and lung parenchyma is crucial for effective gas exchange. The extracellular matrix (ECM) provides the lungs with the elasticity needed for breathing as well as the structural support for resident and infiltrating cells. This dynamic and bioactive network is composed of collagens, fibronectin, laminin, periostin, glycosaminoglycans, hyaluronic acid, and elastin, among others [1]. The twenty eight types of collagens can be categorized into fibril-forming collagens, network-forming collagens, fibril-associated collagens, and transmembrane collagens [2]. In the lung, collagen type I and III are the most abundant fibrillar collagens, while the basement membrane-forming collagen type IV is the major network-forming collagen [3]. The structural organization of the lung ECM importantly relies on collagen organization through collagen crosslinking and interactions with other ECM components [1]. The organization of fibrillar collagens into collagen fibers occurs in the extracellular space. This process is orchestrated in two distinct ways: 1) fiber crosslinking via enzymes such as lysyl oxidases or transglutaminases that are crucial in maintaining collagen stability [1, 2], or 2) fiber organization through interactions with other ECM components [4] such as decorin or biglycan that can significantly impact collagen fibril structure [5–7]. Overall, lung ECM organization affects biomechanical properties such as stiffness and stress relaxation and ultimately cell behavior [8–11].

In the alveolar region of the lung, collagen type I and III are the major load bearing proteins while elastin fibers drive recoil pressure at low strains [12, 13]. At low lung volumes, collagen fibers remain crimpled and wavy but they progressively straighten and stiffen with increasing strains as the lung inflates [12, 13]. Together this fiber realignment imparts the lung with protective non-linear elastic behavior, also known as strain-hardening. Thus, any pathological disruption to collagen organization and crosslinking can severely impact mechanical properties of the lung. In pathological conditions, aberrant strain hardening responses could lead to overly stiffened ECMs at higher lung volumes resulting in impaired compliance and difficulty in breathing. Cell-ECM interactions are vital in regulating processes during development and disease [14], and these interactions in the lung can occur via cell-collagen interactions, among many others. Cells interact with collagens through integrins or mechanoreceptors sensing collagen’s biochemical presence or biomechanical properties (stiffness or prestress of fibers) [1, 15]. Collagen topography can also direct cellular behavior. Studies have separately investigated the influences of the biochemical composition or biomechanical properties of the microenvironment on cell behavior over the last couple of decades [16–24]. Alterations in the ECM organization during the development of different lung diseases and their implications on cell behavior have not been explored thoroughly.

Chronic lung diseases, such as chronic obstructive pulmonary disease (COPD) or fibrotic lung diseases, such as idiopathic pulmonary fibrosis (IPF), are strongly associated with irreversible changes in ECM structure at the disease end-stage [11]. The lung parenchyma of patients with IPF displays excessive deposition and crosslinking of ECM, leading to stiffer and less elastic tissue [3, 25]. On the other hand, patients with COPD often have small airway disease and emphysema as a result of fibrosis around the small airways and destruction of parenchymal ECM respectively [26, 27]. Due to the complex nature of lung ECM, replicating these broad and simultaneous changes using synthetic materials for the development *of in vitro* models is very challenging [28]. However, by using patient-derived lung tissue to generate scaffolds of decellularized ECM, it is possible to study the effects of collagen organizational changes [29]. Importantly, before further investigations of the influence of collagen organization on cell behavior can be advanced, establishing a thorough characterization of collagen organization in healthy and diseased samples derived from lung tissue is key for understanding any baseline changes between these lung states.

Collagen organization has been studied previously in several tissues using various methods. Early efforts focused on investigating collagen organization through the means of (proteomic) content analysis. By analyzing the ECM remodeling enzymes and collagen-interacting proteins, an inferred status of the collagen organization was established [1, 30, 31]. Other methods include measuring total collagen by acid hydrolysis and crosslinking by high-performance liquid chromatography [32, 33]. Parallel to these, imaging-based assays have also been employed to study collagen organization, including staining with Picrosirius Red that provided valuable clues regarding collagen fibrils and their organization [34, 35]. Second harmonic generation microscopy has provided significant opportunities to visualize fibrillar collagens [36, 37]. This powerful and stain-free method relies on the natural light reflecting and refracting properties of the non-centrosymmetric assembly of certain collagen triple helices once these interact with photons in two-photon microscopy. While a few studies have investigated collagen organization in lung tissue as an indicator of lung disease progression or cellular responses [38–40], characterization of collagen organization using combinations of the methods mentioned above in control and diseased lung tissue has not been previously performed.

In the current study, we hypothesized that collagen crosslinking and organization are altered in fibrotic and COPD lungs compared to control. We aimed to provide further insight into the organizational changes in collagen towards understanding the altered tissue responsiveness in chronic lung diseases. To this end, characterization of decellularized lung tissue samples obtained from control donors and patients with COPD or fibrotic lung disease using ultra-high performance liquid chromatography to assess collagen crosslinking and visualization with second harmonic generation microscopy to investigate collagen organization was related to the ECM mechanical properties.

## Methods

### 1. Ethics and subjects

All COPD, one fibrotic and two control lung tissues were obtained at the University Medical Center Groningen (UMCG, The Netherlands) following cancer lung resection surgery or lung transplantation. The tissue was obtained as distal as possible from the resected tumors. Macroscopic examination was performed by a lung pathologist to verify that no tissue abnormalities were present in the obtained lung samples. Experiments in this study were conducted in accordance with the Research Code of the University Medical Centre Groningen (UMCG) stated on https://umcgresearch.org/w/research-code-umcg and national ethical and professional guidelines of the Code of Conduct for Health Research that can be found here https://www.coreon.org/wp-content/uploads/2023/06/Code-of-Conduct-for-Health-Research-2022.pdf (last accessed: April 5, 2026). Archival lung tissue used in this study was exempt from the Medical Research Human Subjects Act in the Netherlands as confirmed by the Medical Ethical Committee of the UMCG. The protocol was approved by the UMCG Central Ethics Review Board for non-WMO studies (study no. 10748) and exempt from consent in compliance with national laws (Dutch Laws: Medical Treatment Agreement Act (WGBO) art 458 / GDPR art 9/ UAVG art 24). All donor material and clinical data were deidentified prior to provision for research purposes. Remaining lung tissue, provided by the University of Michigan (five fibrotic and four control tissues), was deidentified and acquired from deceased donors exempting the study from oversight as deemed by the Institution Review Board of the University of Michigan. The clinical characteristics of lung tissue donors are summarized in **Table 1**.

**Table 1:** Donor characteristics of patients included in this study.

|  | Control (n= 6) | Fibrotic (n= 6) | COPD (n= 6) | p-value |
| --- | --- | --- | --- | --- |
| **Age (median ± range)** | 53.0 (30.0-63.0)* | 53.8 (50.0-67.0) | 56.0 (51.0-61.0) | ns |
| **Sex (Females/Males)** | 2/2* | 0/6 | 2/4 | ns |

*\*Clinical characteristics were unavailable for two donors. Differences between the groups were tested using Kruskal Wallis with Dunn's correction (age) and Chi-square test (sex). A p value <0.05 was considered significant. Ns = not significant.*

### 2. Decellularization

For decellularization, cartilaginous airways and large blood vessels were removed from control, fibrotic and COPD lungs. Decellularization was performed as described previously with slight modifications [41]. Briefly, lung tissue was cut into 1-1.5 cm^3^ pieces and washed thrice with Milli-Q® (3000g) at room temperature. The following treatments were carried out sequentially: Triton X-100 (1%, Sigma-Aldrich, Saint Louis, USA), sodium deoxycholate (2%, Sigma-Aldrich), sodium chloride (1M, Sigma-Aldrich), and DNase (30µg/mL, Sigma-Aldrich) in a MgSO_4_.7H_2_O (1.3mM, Merck, Burlington, USA) and CaCl_2_ (2mM, Merck). Each treatment was carried out at 4°C for 24 hours, except the DNase step which occurred at 37°C, with constant rotation. Between each step, the tissue blocks were washed thrice with Milli-Q® (3000g) at room temperature. The treatments were repeated twice to ensure efficient decellularization. Finally, the decellularized scaffolds were sterilized using 0.18% peracetic acid (Sigma-Aldrich) prepared in 4.8% ethanol for 24hours at 4°C with constant rotation. Afterwards, the material was washed thrice with sterile Dulbecco’s phosphate buffered solution (DPBS) (Gibco, Waltham, MA, United States) and stored in penicillin/streptomycin (1%, Gibco) solution in PBS at 4°C until use. The decellularized ECM (dECM) scaffolds were then either prepared for second harmonic generation microscopy or sectioned for evaluating the degree of crosslinking using chromatography (see below for details).

### 3. Sample preparation

The dECM scaffolds were fixed with 2% paraformaldehyde (Acros Organics, Geel, Belgium) for 30minutes at room temperature. The paraformaldehyde was aspirated and the samples were washed with DPBS. The scaffolds were embedded in 1% UltraPure agarose (Invitrogen, Carlsbad, USA) solution. The agarose-embedded samples were paraffin-embedded and cut into 4 or 15µm thick sections. The 4µm sections were used for standard hematoxylin and eosin staining for visualizing the characteristics of the scaffolds, while 15µm sections were used for crosslinking measurements or for second harmonic generation imaging.

### 4. Hematoxylin and eosin staining

The dECM scaffolds were visualized by standard hematoxylin and eosin staining. Briefly, 4µm scaffold sections were deparaffinized and stained with hematoxylin and eosin using a standard staining protocol. Next, the slides were fixed with 4% paraformaldehyde and mounted with coverslips using Aquatex mounting medium (Merck). The tissues were scanned using the Hamamatsu Nanozoomer 2.0 HT (Hamamatsu Photonic K. K., Japan) at 40x magnification. All the control sections were verified by a lung pathologist (M. Zwager) to confirm the gross normal structure.

### 5. Measuring total collagen and degree of crosslinks

Lung dECM scaffolds were cut into 15µm sections and total collagen content and the degree of crosslinking were assessed in a pooled sample of three serial sections per donor. Samples were deparaffinized, homogenized and then completely hydrolyzed in 6M HCl. Total collagen was measured in the hydrolysates using a sensitive tissue collagen assay (Quickzyme, Leiden, the Netherlands) according to the manufacturer’s instructions. Then the hydrolysates were dried under nitrogen flow, and reconstituted in elution buffer (48007, Chromsystems Instruments & Chemicals GmbH, Gräfelfing, Germany). Pyridinoline crosslinks were analyzed by ultra-performance liquid chromatography (UPLC) (ACQUITY UPLC H-Class, Waters Chromatography B.V., Breda, the Netherlands) equipped with a high-performance liquid chromatography column (48100, Chromsystems) using an isocratic separation method (mobile phase 48001, Chromsystems; flow rate 1.2 mL/min) with subsequent fluorescent detection (EX 290 nm, EM 395 nm; ACQUITY UPLC Fluorescence Detector, Waters). Pyridinoline crosslinks were expressed as crosslinks/collagen (mol/mol).

### 6. Second harmonic generation microscopy

Prior to second harmonic generation imaging, 15µm dECM lung sections were deparaffinized and mounted using Aquatex mounting medium (Merck) and covered with 0.17mm thick coverslips (Epredia, the Netherlands). Second harmonic generation microscopy signals were detected in dECM lung scaffolds using the Stellaris 8 CRS (Leica, Germany) microscope. The excitation wavelength was set at 1031.1nm throughout the experiment and to maximize collagen detection a quarter wave plate was used (Bernhard Halle Nachfl. GmbH, Berlin, Germany). The incident laser power was maintained at 0.3W throughout the experiment. The forward (52.3%) and backward (60.5%) second harmonic generation gain and intensity (60%) were kept constant while imaging different samples. A 40x (HC PL IRAPO 40x/1.1 W) water immersion objective and a numerical aperture 0.9 condenser lens were used for the detection of the forward and backward signals. Kohler illumination was carefully calibrated to ensure optimal detection of the forward signal. The same section of healthy porcine tendon was used to calibrate the microscope before every imaging session. The second harmonic generation signal was detected by a forward and backward detector with a 510/20nm bandpass filter and the autofluorescence was recorded by the epi (E-) coherent anti-Stokes Raman Scattering detector. For every sample, an overview (approximately 3×2mm) of the scaffold was imaged. LASx (Leica) software was used to process and extract the images.

### 7. Image analysis

#### 7.1 Surface area and volume of dECM scaffolds

The estimated surface area of each dECM scaffold was calculated using Fiji [42]. The sections stained with hematoxylin and eosin were digitalized using a slide scanner and visualized using Aperio ImageScope (v12.4.6.5003) and any excessive background or artefacts were removed using Adobe Photoshop (2024). The images were converted to an 8-bit grey scale. The total area of the scaffold was measured by drawing a region of interest (ROI) around the scaffold on the scanned image, and the area of tissue within the scaffold ROI was calculated using a cut-off threshold that enabled selection of only pixels that contained tissue within the scaffold ROI. One threshold setting per group was selected (Control= 0-212, fibrotic= 0-213, COPD= 0-215). Non-tissue spaces were calculated by subtracting total tissue positive pixel area from the total ROI area (equation i). Non-tissue percentage was determined with respect to total ROI area (equation ii). Volume was determined by multiplying the obtained tissue surface area by the 45 (3×15µm) µm which equates to the total thickness of the tissue used in the analyses.

i. *Non* − *tissue* (*pixels*) *area* = *Total area* − *total tissue area*
ii. 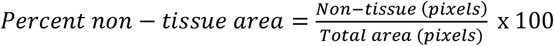
iii. *Volume* = *Total tissue area x* 45µm

For total area (mm^2^), total tissue area (mm^2^), and volume (µg/mm^3^) of scaffolds mean ± standard deviations have been reported.

#### 7.2 Second harmonic generation image analyses

Second harmonic generation images were used to characterize gross ECM organization and collagen modifications as illustrated in **Figure 1**. Each image was thresholded and binarized to identify pixels corresponding to the decellularized tissue. Next, holes within the tissue were identified as background pixels surrounded by the matrix. Identified holes were filtered by minimum area and maximum aspect ratio. This selection was made based on average shape and alveoli size reported in literature for control, fibrotic and COPD lungs [43–45]. As filtering thresholds, a lower hole-diameter of 100µm and a maximum aspect ratio of 5 were applied to discard artifacts or technical biases introduced during tissue sectioning and embedding. To quantify the ratio of holes versus tissue per image, the total area of the selected holes was divided by the total area of the tissue. Next, collagen modifications were assessed by investigating forward to backward signal ratios that were generated per pixel. To compare collagen organization among control and diseased models, median forward to backward ratios that were derived from a 15-pixel wide ring around each identified hole were assessed. Data from all donors within each group were pooled together to study forward to backward signal versus hole area tendencies and were sorted by hole area. Mean ± standard deviations were reported for area of holes relative to tissue area, median area of holes and F/B intensity for each group.

**Figure 1:**
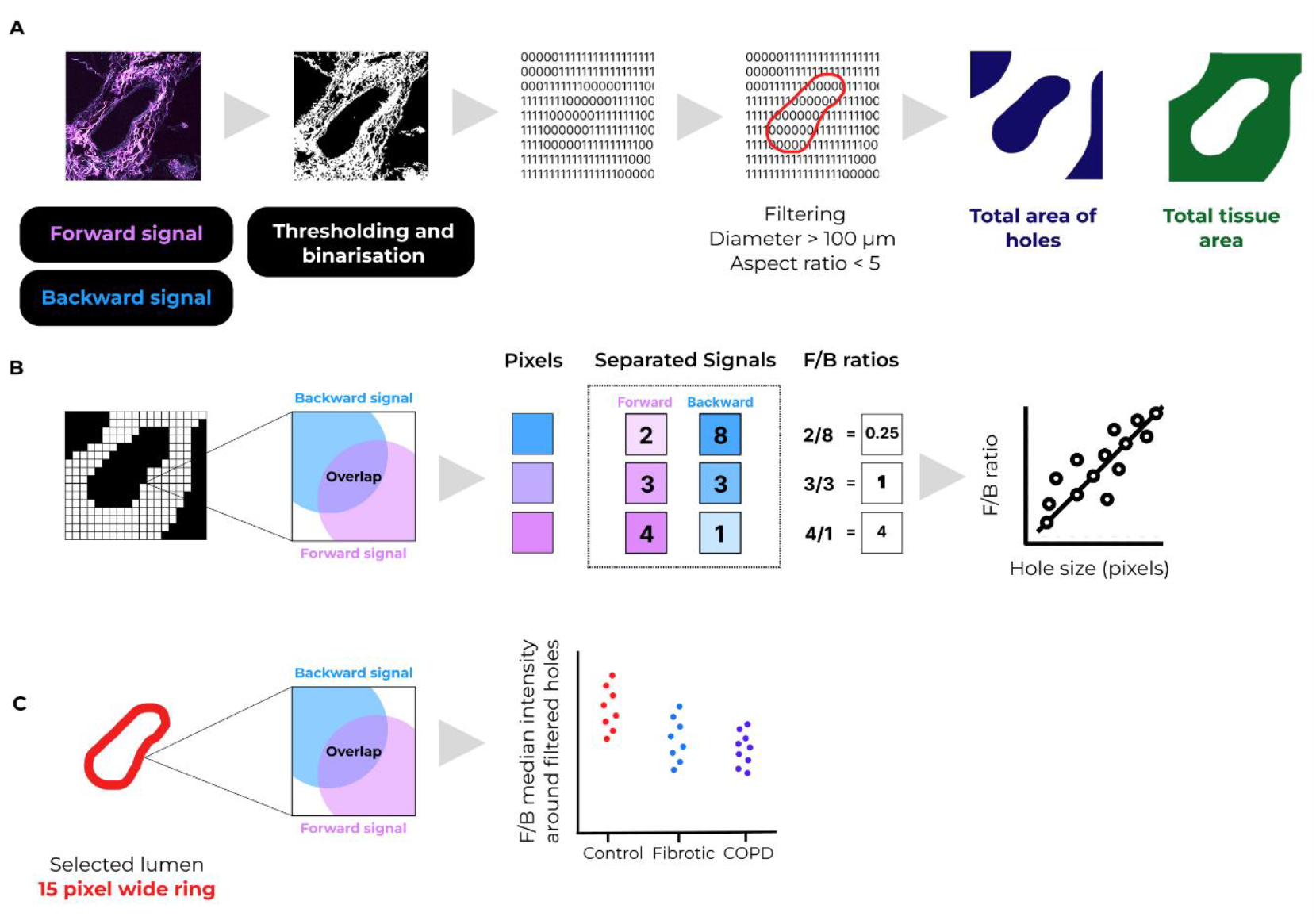
Analysis workflow for second harmonic generation images. A) Second harmonic generation images were converted to 8-bit images, thresholded, and binarized to identify total area of pixels belonging to tissue. Holes within the tissue were identified as background pixels surrounded by continuous matrix pixels. Holes that had diameters above 100µm or aspect ratio less than 5 were selected for further analysis. Total hole area was calculated relative to total tissue area. B) Collagen modifications were assessed by generating the forward to backward ratio per pixel for the whole image and plotted for increasing hole size. C) The three groups were compared by measuring forward to backward ratios within a 15-pixel ring around each identified hole. Six donors were analyzed per group.

### 8. Strain hardening

Mechanical data obtained from low load compression testing of lung tissue in a previous study were [9] reanalyzed to evaluate strain hardening of fresh human lung tissue across the lung disease groups of the study. Stress relaxation was previously measured in control (n= 6), fibrotic (n=6) and COPD (n= 14) lung tissues using a low load compression tester. Young’s modulus for selected donors was presented in the previous study, while in the current studied all donors were included. Lung tissues obtained from COPD patients were from different GOLD (Global Initiative for Chronic Obstructive Lung Disease) stages of the disease including stage I (n = 4), stage II/III (n=2) and stage IV (n=8). Lung tissues were subjected to 20% strain, which was held constant for 200 seconds to measure stress relaxation. Young’s moduli were calculated at 5% and 10% strain by fitting stress versus strain curves on five datapoints in the vicinity of 5% strain and 10% strain. Strain hardening was further quantified as the ratio of Young’s modulus at 10% strain to that at 5% strain. Three measurements were recorded per donor which were averaged for further analysis.

### 9. Statistical Analyses

Prior to comparison, data for each parameter measured were tested for normality using QQ plots **(Figure S1)**. Age and sex distribution between different groups were compared using Kruskal Wallis with Dunn’s correction and Chi-square test, respectively. Sample parameters, including area, volume, degree of crosslinking, strain hardening and relative and median area of holes were compared between control, fibrotic, and COPD using one-way ANOVA with Tukey’s or Dunnett’s multiple test correction. Associations between the degree of crosslinking and total collagen or tissue or non-tissue area were evaluated using linear regression analysis. Non-linear regression analysis identified associations between forward to backward ratios and hole sizes, while Kruskal-Wallis with Dunn’s correction was used to compare median forward to backward ratios and strain hardening among control, fibrotic, and COPD donors. For all analyses, a p-value < 0.05 was considered significant. GraphPad Prism (10.6.1 (892)), IBM SPSS 28.0.1.0(142), and R studio (2023.03.0 + 386) were used for statistical analyses.

## Results

### 1. Decellularized control, fibrotic, and COPD dECM scaffolds resembled characteristics of native lung tissue

In order to assess whether dECM scaffolds retain anatomical features of native lung tissue, dECM scaffolds derived from control, fibrotic, and COPD lung tissue were visualized by staining with hematoxylin and eosin **(Figure 2)**. Qualitatively, control dECM scaffolds had the appearance expected of normal lung tissue, with intact alveoli, airway walls, and blood vessels all visible. Fibrotic dECM scaffolds had more ECM deposition around the airway and blood vessel walls and within the alveoli, compared to control and COPD scaffolds. In contrast, COPD scaffolds had thickened airway walls with damaged alveoli and loss of parenchymal tissue.

**Figure 2:**
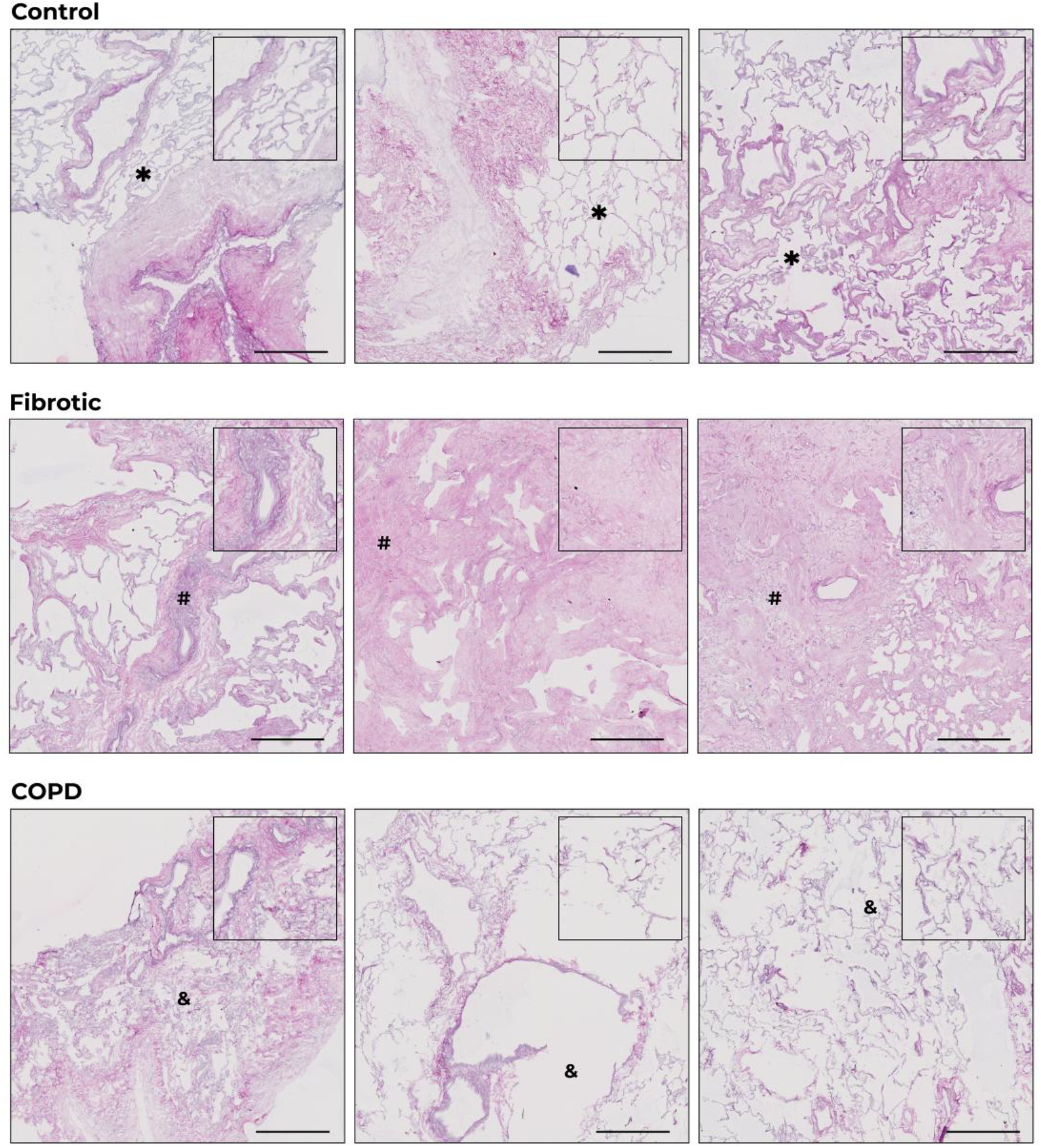
Representative regions of control, fibrotic, and COPD dECM lung scaffolds stained with hematoxylin and eosin. Representative regions of control (n=6) fibrotic (n=6), and COPD (n=6) decellularized scaffolds are shown, taken from lung tissue sections stained using hematoxylin and eosin and digitally captured using a whole tissue section scanner. Areas depicting typical features of control (normal and intact alveoli, airway, and blood vessel walls), fibrotic (excessive deposition of ECM in parenchyma), and COPD (thickened airway walls and damaged alveoli with enlarged airspaces) tissue are highlighted in the higher magnification inserts and marked by *, # and & respectively. Scale bars=500µm. COPD: chronic obstructive pulmonary disease; dECM: decellularized extracellular matrix.

### 2. Fibrotic dECM scaffolds had larger tissue surface area and a lower percentage of non-tissue area than control and COPD dECM scaffolds

To quantify the visual differences between control, fibrotic, and COPD dECM scaffolds, tissue area and estimated volume of the scaffolds were calculated. Fibrotic scaffolds had the largest tissue area (101.6 ± 30.84 mm^2^), followed by control (64.68 ± 16.06 mm^2^), while COPD scaffolds had the smallest tissue area (38.98 ± 8.86 mm^2^) **(Figure 3A)**. As expected, total tissue volume followed the same pattern as the tissue area, with fibrotic scaffolds having the highest volume (4.57 ± 1.39 mm^3^), followed by control (2.91 ± 0.72 mm^3^) scaffolds, and COPD (1.75 ± 0.40 mm^3^) **(Figure S2)**. Subsequently, the percentage of non-tissue areas within the scaffolds (airway or blood vessel lumens or enlarged airspaces) was calculated. COPD scaffolds had a higher percentage of non-tissue areas (69.29 ± 11.05%) compared to control (61.84 ± 9.83%) or fibrotic (49.39 ± 6.72%) scaffolds **(Figure 3B)**.

**Figure 3:**
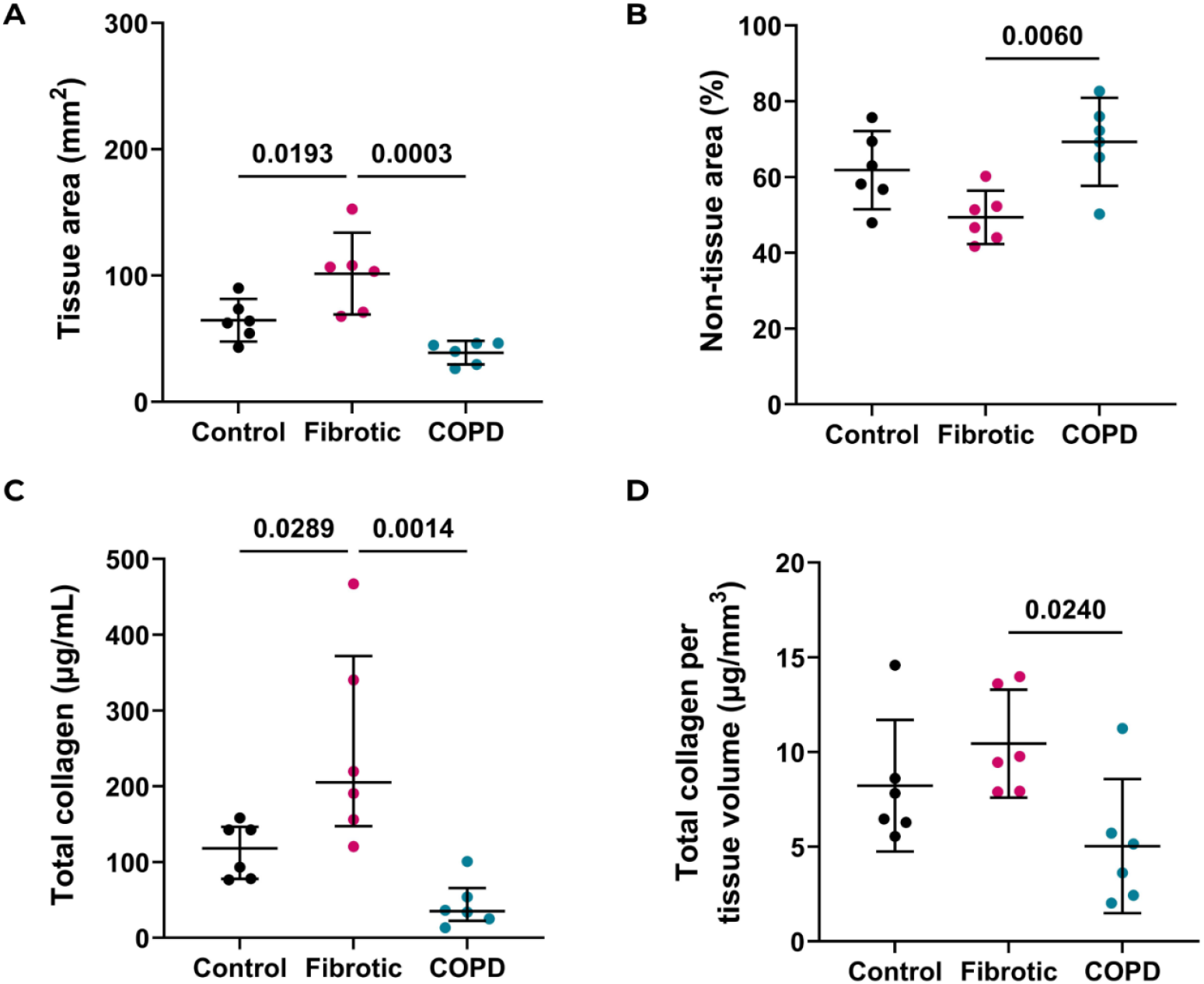
Total tissue and non-tissue area and amount of collagen in dECM control, fibrotic and COPD lung scaffolds. A) Total tissue area and B) percentage non-tissue area of formalin-fixed paraffin-embedded decellularized lung scaffolds from control (n=6), fibrotic (n=6), and COPD (n=6) were calculated using image analysis of hematoxylin and eosin-stained sections. C) Absolute amount of total collagen was quantified using a sensitive tissue collagen assay (Quickzyme) D) and was normalized to tissue volume. Mean with 95% CIs are plotted and each dot represents a dECM scaffold from a different donor. One-way ANOVA with Tukey’s correction was used to test the differences between the groups. A p-value <0.05 was considered significant. COPD: chronic obstructive pulmonary disease; dECM: decellularized extracellular matrix.

### 3. COPD dECM scaffolds had lower amounts of total collagen but a greater degree of crosslinking per mole collagen

We examined the total amounts of collagen in control, fibrotic, and COPD dECM scaffolds **(Figure 3C)** and normalized these to tissue volume **(Figure 3D)**. COPD (5.03 ± 3.37 µg/mm^3^) scaffolds had less total collagen compared to control or fibrotic scaffolds, while no difference was noted in total collagen content between fibrotic (10.44 ± 2.71µg/mm^3^) and control (8.22 ± 3.31µg/mm^3^) scaffolds.

The degree of crosslinking per mole of collagen was highest in COPD scaffolds, with an average of 0.22 ± 0.04 pyridinoline crosslinks per mole of collagen compared to 0.12 ± 0.02 in fibrotic scaffolds and 0.08 ± 0.05 in control scaffolds **(Figure 4A)**. Moreover, the degree of crosslinking in the different groups was negatively associated with the amount of total collagen normalized per tissue volume (p= 0.033) **(Figure 4B)**.

**Figure 4:**
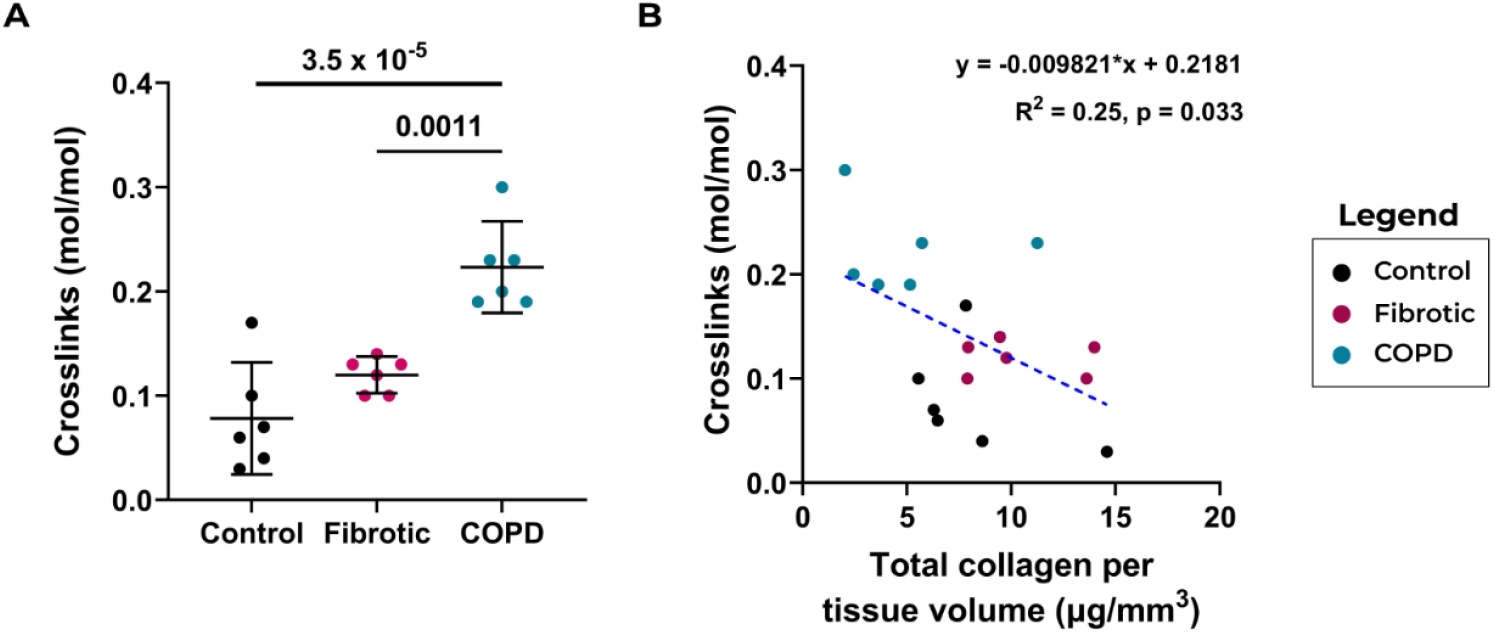
Quantification of collagen crosslinks in control, fibrotic and COPD dECM lung scaffolds. A) Pyridinoline crosslinks were measured using ultra-pure liquid chromatography in formalin-fixed paraffin-embedded decellularized lung scaffolds from control (n=6), fibrotic (n=6), and COPD (n=6). B) Linear regression models were used to evaluate the association between total collagen normalized to tissue volume and the degree of crosslinking. Mean with 95% CIs have been plotted and each dot represents a dECM scaffold from a different donor. One-way ANOVA with Tukey’s correction was used to test the differences between the groups. A p-value <0.05 was considered significant. COPD: chronic obstructive pulmonary disease; dECM: decellularized extracellular matrix.

### 4. Mature crosslinked collagens were localized to airway and blood vessel walls in control and COPD dECM scaffolds but were spread throughout fibrotic dECM scaffolds

Having observed a difference in the degree of collagen crosslinking, collagen organization in control, fibrotic, and COPD dECM scaffolds was studied using second harmonic generation microscopy. The non-centrosymmetric structure of fibrillar collagen produces an endogenous second harmonic signal without exogenous labels. Upon excitation of collagen fibers, light is emitted with half the wavelength and the direction of propagation of light is directed by the maturity (organization) of the collagen fibers. A signal captured at the forward detector reflects the presence of mature crosslinked and organized collagen, while immature disorganized fibrils with less crosslinking scatter light to the backward detector [46]. **Figure 5A** exemplifies varied regions of control, fibrotic, and COPD scaffolds, while overviews of all the scaffolds imaged in the current study can be seen in **Figure S3**. Control, fibrotic, and COPD dECM scaffolds had distinct localizations of the forward (mature collagens) and backward (immature collagens) signals. Control scaffolds were characterized by mature crosslinked collagens concentrated within the vicinity of the airway and blood vessel walls accompanied by mainly immature collagens in the parenchyma **(Figure 5B)**. In fibrotic scaffolds, mature crosslinked collagens were detected throughout the scaffolds, irrespective of the compartment, whereas immature collagens were noted predominantly in the parenchyma. In COPD dECM scaffolds, mature crosslinked collagens were concentrated around the airway and blood vessel walls. As airway walls were thickened in COPD donors, the area of mature collagens around airway walls appeared larger than controls. The remaining alveolar walls in COPD tissue were composed of immature collagens. In summary, mature collagens were predominantly a feature around airways and blood vessel walls in dECM lung scaffolds of control, fibrotic, and COPD groups. In fibrotic scaffolds, mature collagens were also present within the parenchymal region and immature collagens were mainly detected in the parenchyma of control and COPD scaffolds **(Figure 5B)**.

**Figure 5:**
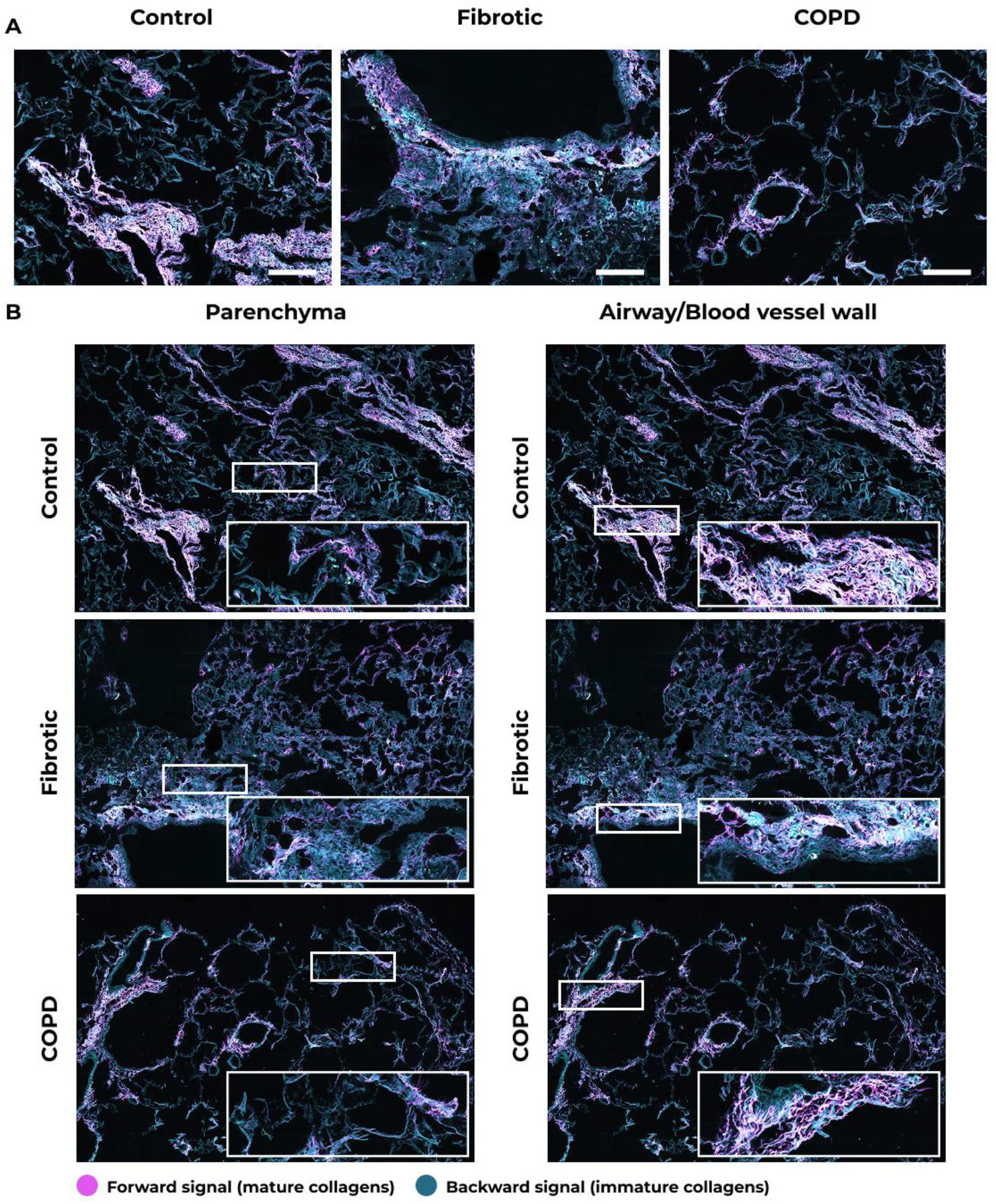
Differential localization of mature and immature collagens in control, fibrotic, and COPD dECM scaffolds. Second harmonic generation microscopy was used to study collagen organization in control (n=6), fibrotic (n=6) and COPD (n=6) decellularized lung ECM scaffolds. Mature collagens are highlighted in magenta and immature collagens in cyan. A) High magnification representative images of control, fibrotic, and COPD dECM scaffolds. Scale bar=250 µm. B) Magnified regions (parenchyma or airway/blood vessel walls) highlighting the differences in localization of mature organized (forward signal in magenta) and immature disorganized (backward signal in cyan) collagens between groups. COPD: chronic obstructive pulmonary disease; dECM: decellularized extracellular matrix.

### 5. Fibrotic tissue had higher median forward to backward signal intensity ratio around defined holes

Next, the intensity of forward and backward signals across all pixels of each image were examined. Averaging the areas or intensities of forward and backward signals of all pixels per tissue from each donor **(Figure S4E-H)** failed to fully capture the differences between groups, as illustrated in **Figure 5**, and obscured the observed nuances in the images. To overcome this disparity, collagen organization was investigated specifically within a 15-pixel wide ring around identified holes in each dECM scaffold image. The area of holes relative to tissue (control= 0.54 ± 0.15, fibrotic= 0.39 ± 0.18, COPD= 0.58 ± 0.17) and median area of holes (control= 2073 ± 455.2, fibrotic= 1825 ± 289, COPD=1889 ± 173.2 pixel^2^) was similar in all groups **(Figure 6A** and **B)**. Subsequently, the relationship between the forward to backward signal ratio (F/B ratio) and hole size was analyzed across groups **(Figure 6C-E)**. While the F/B ratio remained consistent across hole size in control and COPD donors, a trend towards increasing F/B ratio with increasing hole size was noted in fibrotic tissue. These intensities were then compared among control and diseased groups **(Figure 6F)**. Fibrotic tissue had higher median F/B ratios (0.84 ± 0.74) around the defined rings as compared to both control (0.39 ± 0.45) and COPD donors (0.35 ± 0.41).

**Figure 6:**
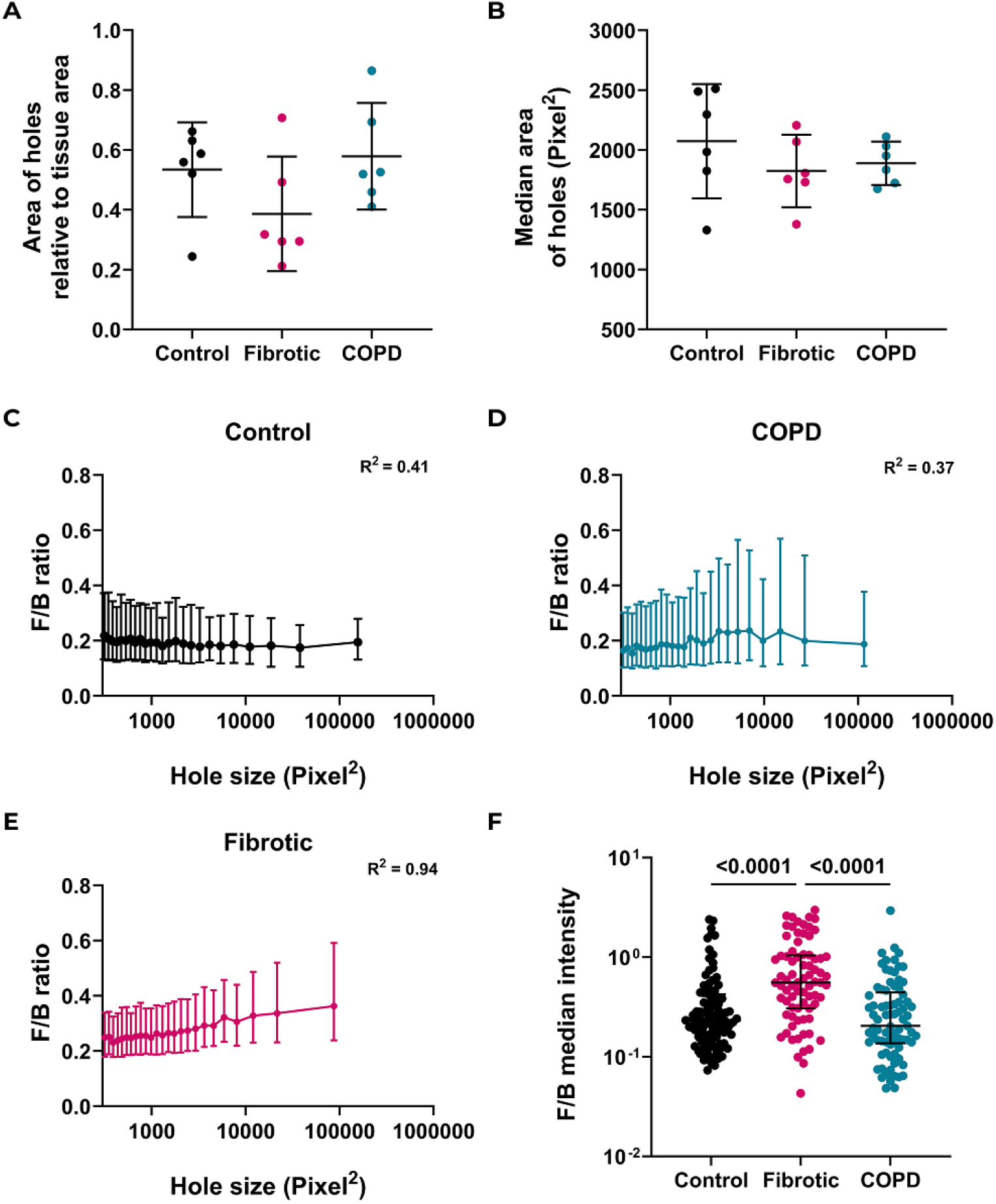
Quantifying distribution of forward and backward signal across tissues. A) Area of defined holes relative to tissue area and B) Median area of holes in control (n=6), fibrotic (n=6) and COPD (n=6, stage IV) dECM scaffolds. Means with 95% CI were plotted and compared using One way ANOVA with Dunnett’s correction. The distribution of F/B ratios across increasing hole sizes were depicted in C) through E) for each group (n=25 per group). Mean ± standard error was plotted for each group. F) Comparison of median F/B intensities for each identified hole and surrounding rings in control (n=99), fibrotic (n=77) and COPD (n=81) dECM scaffolds. Median with the interquartile range was plotted, and Kruskal Wallis with Dunn’s correction was used to test differences. A p-value <0.05 was considered significant. COPD: chronic obstructive pulmonary disease; dECM: decellularized extracellular matrix.

### 6. Fibrotic human lungs exhibit higher strain hardening compared to control and COPD

Our data revealed that COPD dECM lung scaffolds had a greater degree of crosslinking per mole collagen while fibrotic dECM lung scaffolds had more overall collagen mass weight compared to controls. To assess how increased crosslinking or collagen amount affects lung tissue mechanics in disease, we reanalyzed our previously published lung tissue stress relaxation data [9] to evaluate the strain hardening properties of these lung tissues across the disease groups **(Figure 7)**. Strain hardening is driven by the alignment, stretching or tightening of intra and extracellular fiber networks including actin, fibrin, and collagen resulting in a network that appears to be stiffer when probed at larger strains. As seen in **Figure 7**, fibrotic lungs had higher levels of strain hardening compared to control and COPD donors, indicating an increasing resistance to deformation with increasing strain. Strain hardening in **Figure 7** has only been reported for stage IV COPD donors. A comparison between strain hardening in different COPD stages can be found in **Figure S5**. No differences were noted among different GOLD stages of COPD.

**Figure 7:**
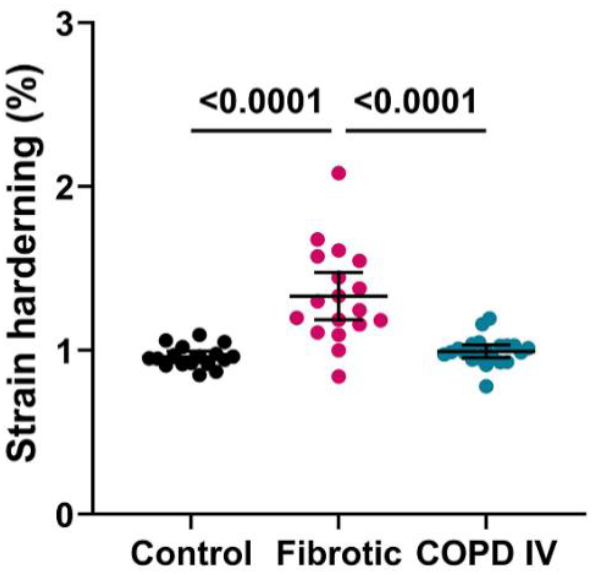
Strain hardening of human lung tissue. Stress relaxation curves obtained from low load compression testing were used to evaluate strain hardening of control (n=6), fibrotic (n=6) and COPD (n=8) human lung tissue and were plotted in triplicates per donor. Mean with 95% CI was plotted and One-way ANOVA with Dunnett’s correction was used to evaluate differences between groups. A p-value <0.05 was considered significant. COPD: chronic obstructive pulmonary disease; dECM: decellularized extracellular matrix.

## Discussion

In the current study, collagen crosslinking and organization were studied in decellularized scaffolds of control, fibrotic, and COPD lungs. We found that control, fibrotic, and COPD dECM scaffolds are representative of native lung tissue in terms of morphological features. Differences in the amount of tissue and non-tissue areas between groups, as expected from the pathological process, were also conserved in diseased scaffolds. Collagen content was higher in fibrotic than both control and COPD scaffolds. However, COPD scaffolds had a higher degree of crosslinking per mole of collagen than the fibrotic or control scaffolds. With respect to organization, fibrotic scaffolds were characterized by mature collagens throughout the scaffold irrespective of the anatomical compartments, while COPD scaffolds displayed mature collagens distributed distinctly in airway and blood vessel walls and immature collagens predominantly in the parenchymal regions. Differences in collagen crosslinking and organization were partially reflected in the mechanical behavior of diseased lungs particularly strain hardening. Fibrotic tissue exhibited higher strain hardening, indicating these tissues exhibited stiffening behavior with increasing strain, whereas the increased crosslinking in COPD scaffolds appeared to partially attenuate this capacity in these tissues.

Distinctive structural features of the control, fibrotic, and COPD lungs were conserved in the respective scaffolds as observed using hematoxylin and eosin staining and second harmonic generation imaging. Fibrotic scaffolds had larger tissue areas and higher amounts of total collagen consistent with higher amounts of deposited collagens reported previously in various studies [47]. In COPD scaffolds, mainly enlarged alveoli contributed to higher non-tissue areas. Additionally, COPD scaffolds had lower amounts of total collagen. Contrary to our results, Eurlings *et al*. (2014), using image analysis, observed an increase in the percentage area of collagen in the alveolar and airway walls of native COPD lungs associated with COPD severity [48]. This disparity in results among studies might be due to the difference in detection techniques, types of quantified collagens and elimination of immature collagens during decellularization resulting in lower total collagen in our samples.

COPD dECM scaffolds had a higher degree of pyridinoline (mature) crosslinking per mole of collagen, than either control or fibrotic scaffolds. This implied that the lung tissue that remains following emphysematous destruction is highly crosslinked. We speculate that fibrotic small airway walls in COPD scaffolds are the main source of the collagen with the higher degree of crosslinking. Small airway disease begins early in COPD disease development, before the manifestation of other symptoms, suggesting prolonged periods for collagen crosslinking within small airways [49–52]. Excessive crosslinking of collagen fibers can confer resistance against degradation mechanisms and provide mechanical reinforcement to small airways to prevent extreme stretch due to loss of alveolar attachments [15]. This is supported by *in vitro* studies where collagen matrices with fewer pyridinoline crosslinks were more susceptible to degradation by matrix metalloproteinase 1 than matrices enriched for pyridinoline crosslinks [53]. As several different types of crosslinks can be present in organized and disorganized collagens, results from the current study could be further extended by measuring different mature and immature crosslinks compartmentally in native control, fibrotic, and COPD lung tissues. However, assays enabling measurements of different types of crosslinks are currently limited.

Second harmonic generation microscopy highlighted differences in collagen organization between control, fibrotic, or COPD scaffolds. Differential localization of forward (mature collagen) and backward (immature collagen) signals in the scaffolds were present that matched the pathological characteristics of fibrotic and COPD lungs. On averaging the overall intensity of forward or backward signals per donor, no differences were noted between groups. To nuance the interrogation of the spatial distribution of the collagen organizational patterns, collagen organization was quantified within a 15-pixel ring surrounding holes within the scaffolds. Fibrotic tissue had higher F/B median intensity within these zones compared to control and COPD groups which was in concordance with a study that demonstrated increased ratio of organized to disorganized collagen in IPF parenchymal tissue but did not investigate IPF airways [54]. A higher F/B ratio could indicate a combination of higher forward signal and lower backward signal implying overall more mature organized collagens around defined holes in fibrotic tissue in comparison to control and COPD donors. In another study, a decreased ratio of organized to disorganized collagen has been reported in COPD airways [38]. Our results on decellularized lung ECM build on the results obtained using native lung tissue [55] and illustrate complex spatial distribution of collagen organization in dECM scaffolds.

Lung tissue typically exhibits nonlinear strain-stiffening behavior arising from the mechanical interactions between collagen and elastin fibers, and other ECM components. Elastin governs low-strain response and provides compliance, while crimped collagen fibers progressively straighten under increasing strain, causing a nonlinear rise in stiffness [13]. Fibrotic lungs exhibited higher strain-hardening than the other two tissue groups, consistent with increased collagen content and more organized collagen architecture. From the mechanical perspective, COPD lungs showed stress-strain behavior comparable to controls. Nevertheless, from the organization perspective, COPD lungs had less total weight of collagen than controls, but more crosslinked collagen (mol/mol) than controls. Together, these findings highlight the compensatory mechanical role between scaffolding proteins and their crosslinkers, that make it difficult to predict not only the stiffness, but also the strain-hardening behavior of biological tissue. Using multiphoton microscopy, Deng *et al*. (2026) showed that fibrotic human precision cut lung slices (hPCLS) exhibit increased collagen content and higher fractal dimension, reflecting a denser, more space-filling collagen network. In contrast, COPD tissue showed reduced collagen content and lower fractal dimension, suggesting a disrupted and less connected ECM despite thicker fibers. These structural differences corresponded to decreased stiffness in COPD and increased stiffness in fibrotic tissue and are consistent with our findings [56]. Overall, our findings suggest that collagen content and organization are primary determinants of strain hardening in the lungs. These opposing strain hardening responses in fibrotic and COPD scaffolds are reflected in pressure-volume characteristics-where fibrotic lungs exhibit reduced compliance and restrictive physiology, and COPD lungs have a higher compliance associated with air trapping and airflow limitation.

Collagen organization is regulated by enzymes such as lysyl oxidases (LOX) and transglutaminases and via interactions with proteoglycans and glycoproteins. Disruption of these processes is often linked to disease, and has been suggested in COPD and fibrosis in the lungs. Recent studies showed increased LOX-like protein 1 (LOXL1) and decreased LOX expression in COPD small airways, with transglutaminase 2 levels also elevated associated with disease severity [57, 58]. In fibrotic lungs, LOXL1, LOXL2, LOXL3, and LOXL4 were elevated, and inhibition of lysyl oxidase activity reduced transforming growth factor β-induced stiffening in fibrotic matrices [54, 59]. Beyond the lung, serological levels of LOXL2 in IPF were associated with disease progression [60]. Additionally, IPF fibroblasts exhibited higher expression of transglutaminase 2 expression compared to controls [10]. Selective inhibition of lysyl oxidase proteins in healthy and diseased lungs would highlight the relative contribution of each enzyme. Increased levels of lysyl oxidases and transglutaminases could contribute to a higher degree of crosslinking in COPD and fibrotic lungs compared to controls that we report in this study.

Altered levels of proteoglycans in disease may impact the collagen organization observable through second harmonic generation microscopy. In addition to abnormal levels of lysyl oxidase family proteins, the deposition of proteoglycans and glycoproteins like lumican, decorin, and fibulin 2 is reduced in COPD, while fibrotic foci have increased versican deposition [61–64]. Moreover, in different *in vitro* models of IPF (such as decellularized scaffolds or cell-derived ECM) higher expression and secretion of decorin in IPF and control fibroblasts was noted [65–67]. Proteoglycans such as decorin, lumican and biglycan control collagen fiber diameter and density [7, 68]. Thus, further characterization of collagen fiber length, density, and diameter may reveal alterations in the collagen fiber structural assembly between control, fibrotic, and COPD tissue.

The inverse relationship between total collagen content and collagen crosslinking in our study can initially be seen as counterintuitive. In native IPF lung tissue, we observed a positive association between total collagen amount and mature collagen crosslinking [55]. However, another study noted higher amounts of crosslinks in IPF lung tissue compared to controls without a significant increase in total collagen [59]. Moreover, *in vitro* studies have reported that both the timing and type of anti-fibrotic supplementation significantly influence the composition and maturation of collagen crosslinks [69, 70]. We were unable to detect a difference in the degree of crosslinking between fibrotic and control dECM scaffolds. This finding is not consistent with our results using native lung samples [55] and may be caused by the small number of fibrotic tissues included in our study, decellularization processes, and/or the heterogeneity of the degree of crosslinking detected in the control scaffolds. An increase in crosslinking is often accompanied by increased stiffness, as noted in IPF lung tissue and *in vitro* models [59, 71, 72]. In general, whole IPF lungs are stiffer than control and COPD lungs [9]. Compartmentally though, COPD airway walls are postulated to have higher stiffness, while emphysematous tissue is reportedly softer than its control counterparts [73, 74]. As a step ahead, analyzing stiffness per compartment in dECM scaffolds would help validate our findings.

To our knowledge this is the first study to report collagen crosslinking and organization in dECM scaffolds, revealing stark differences in fibrotic and COPD lungs versus controls. The impact of collagen crosslinking may vary by disease. In COPD, heightened crosslinking could increase airway wall stiffness, causing airflow obstruction, and breathing dysfunction in patients. Although small airways are lost as COPD advances, a higher degree of crosslinking may protect against protease-induced degradation and overstretching of small airways. Investigating the degree of crosslinking and organization in early COPD could explain why some airways are retained and become fibrotic while others are lost. On the contrary, the abundance of total collagen and presence of mature collagens in fibrotic tissue confers increased stiffness and resistance to deformation with increasing strain disrupting the normal functioning of the lung. There are some promising anti-fibrotic candidates, such as GB2064, PX-5505 and ZED1227, that target LOX and transglutaminase activity, but further validation is necessary [75, 76]. Simtuzumab, an inhibitor of LOXL2 activity, prevented and reversed fibrosis in murine models; however, it failed to improve survival in IPF patients [77]. Notably, targeted drug delivery of anti-fibrotic drugs is crucial as inhibition of pan-LOX activity had deleterious effects such as acceleration of elastin-mediated emphysema development [78].

In summary, identifying collagen crosslinking and organization patterns is key for the development of effective and targeted delivery of therapeutics to patients affected by chronic lung diseases. It is important to recognize that investigating collagen content alone may not fully explain the collagen-mediated pathology of a disease and characterization of crosslinking and organization are essential.

## Author Contributions

M.M.J., M.N., and J.K.B. conceived and designed the study. M.M.J., M.N., M.C.M., R. H., T.K., K.S., T.B., and N.G. performed the experiments, collected the data, and conducted the primary data analysis. M.C.M., R. H., T.K., K.S., T.B., M.Z., and N.G. contributed to experimental design, methodology, data interpretation, and critical discussion of the results. M.M.J. and M.N. drafted the manuscript. J.K.B. supervised the project and provided intellectual input throughout the study. M.M.J., M.N., M.C.M, R. H., T.B., M.Z., I.H.H., S.D.P., B.N.M., N.G., and J.K.B. revised the manuscript. All authors reviewed and edited the manuscript, approved the final version, and agree to be accountable for the work.

## Acknowledgments

Authors thank Dr Roderick de Hilster for his stress relaxation measurements and Mr. Albano Tosato for assistance in preparation of Figure 1.

## Data availability

The raw data used to generate the findings of this study are available from the corresponding authors upon reasonable request.

