## Supplemental material for "Differential collagen crosslinking and network organization creates distinct tissue remodeling patterns in fibrosis and COPD"

Janette K. Burgess

University Medical Center Groningen

Department of Pathology and Medical Biology

Hanzeplein 1 [IPC EA11]

9713 GZ Groningen

The Netherlands

### Supplementary material

### Supplementary figures

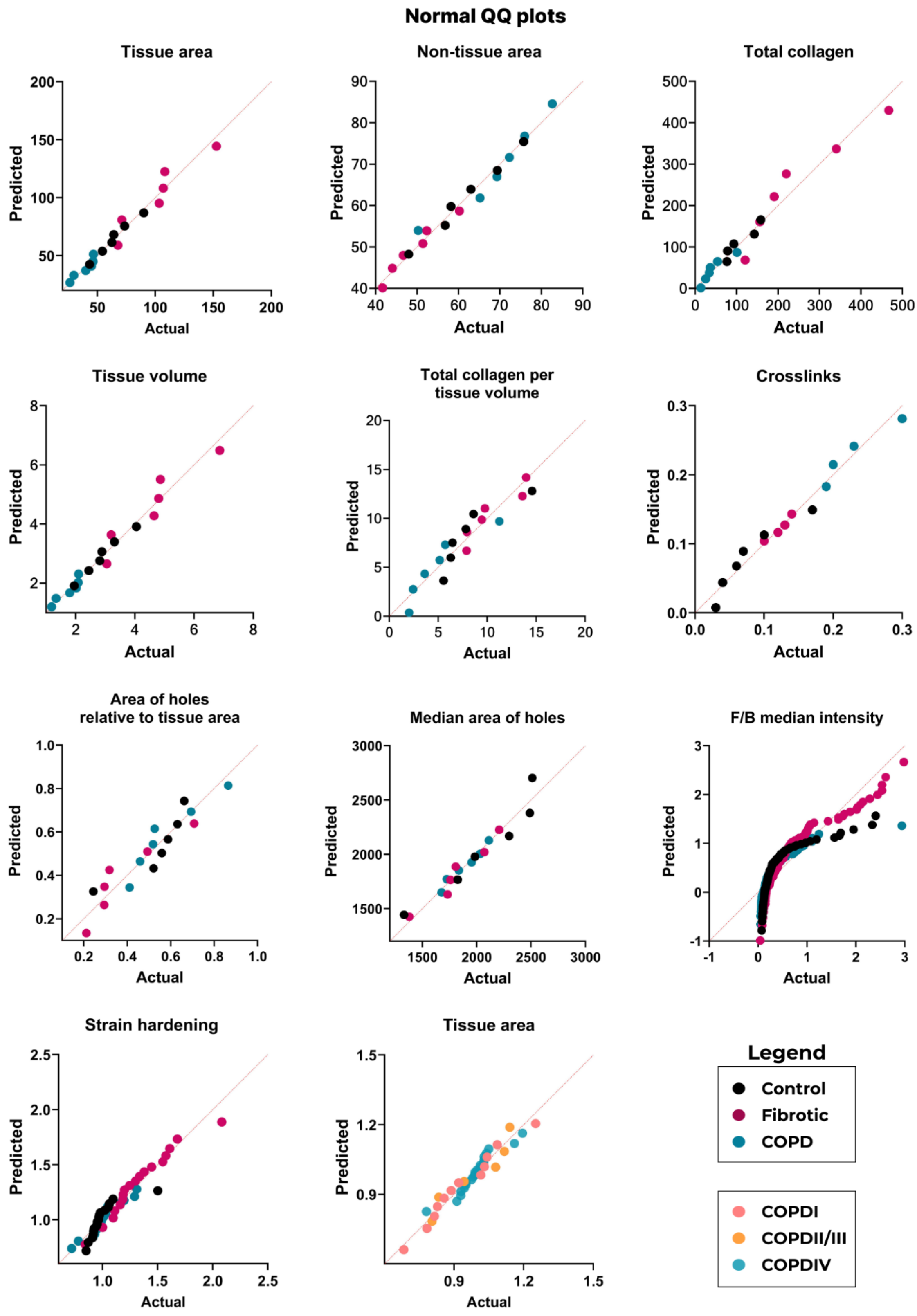

**Figure S1: Distribution of data was determined using QQ plots.** QQ plots were used to visualize the distribution of data for deciding parametric versus non-parametric statistical testing.

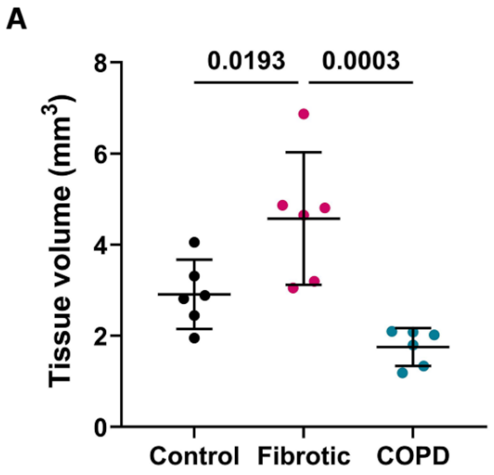


**Figure S2: Volume of control, fibrotic, and COPD dECM scaffolds**. A) Tissue volumes of control (n=6), fibrotic (n=6), and COPD (n=6) were calculated using the tissue area (determined by hematoxylin and eosin staining) multiplied by the thickness of the sections included for analyses. Graphs were plotted using mean with 95% CI and each dot represents a donor. One-way ANOVA with Tukey's correction was used to test the differences between the groups. A p-value <0.05 was considered significant. COPD: chronic obstructive pulmonary disease; dECM: decellularized extracellular matrix.

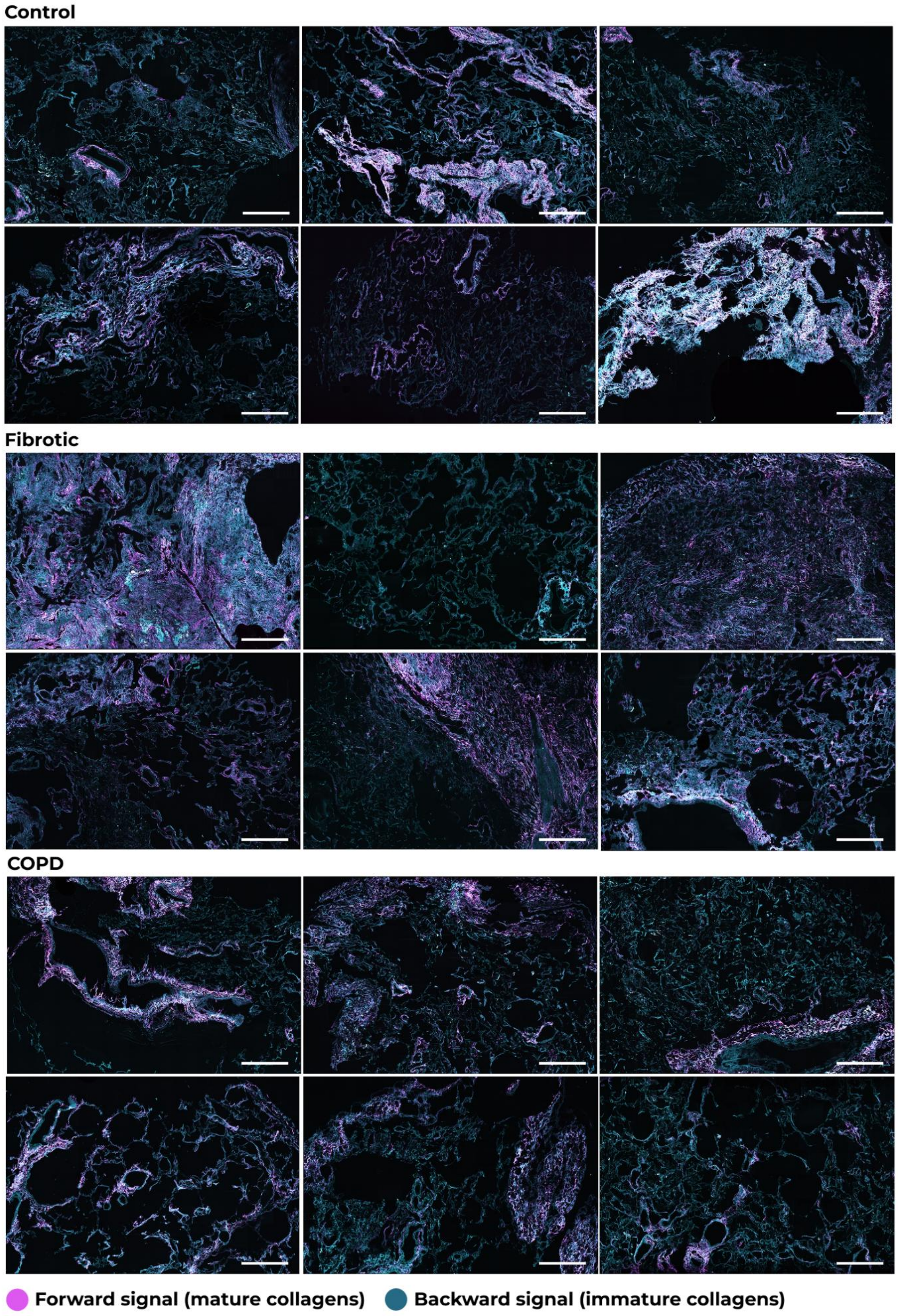


**Figure S3: Overview images for visualization of collagen organization in control, fibrotic and COPD dECM scaffolds.** Overviews of control (n=6), fibrotic (n=6), and COPD (n=6) decellularized scaffolds highlighting mature collagens (magenta) or immature collagens (cyan) imaged using second harmonic generation microscopy. Scale bar=500µm. COPD: chronic obstructive pulmonary disease; dECM: decellularized extracellular matrix.

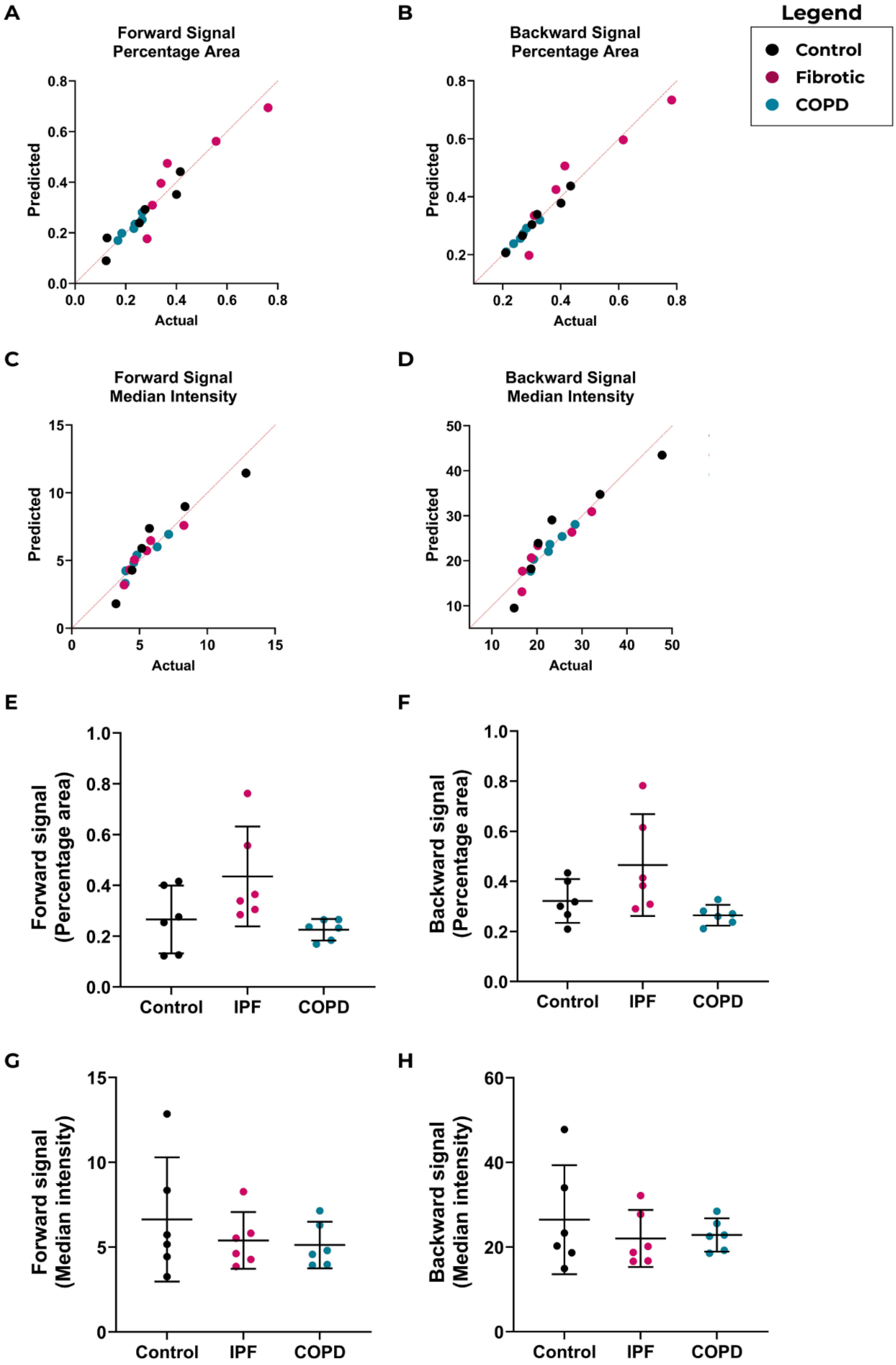
A
Forward Signal Percentage Area
Predicted
Actual
B
Backward Signal Percentage Area
Predicted
Actual
Legend
Control
Fibrotic
COPD
C
Forward Signal Median Intensity
Predicted
Actual
D
Backward Signal Median Intensity
Predicted
Actual
E
Forward signal (Percentage area)
Control
IPF
COPD
F
Backward signal (Percentage area)
Control
IPF
COPD
G
Forward signal (Median intensity)
Control
IPF
COPD
H
Backward signal (Median intensity)
Control
IPF
COPD

**Figure S4: Distribution plots and overall percentage area and median intensity of forward and backward signal**. A) through D) Distribution of data was studied using QQ plots to determine the use of parametric or non-parametric tests. E) through H) Average percentage area and median intensity of forward and backward signal obtained from the overview images for all pixels. Mean with 95% CI was plotted and One-way ANOVA with Dunnett's correction was used to evaluate differences between groups. A p-value <0.05 was considered significant. COPD: chronic obstructive pulmonary disease; dECM: decellularized extracellular matrix.

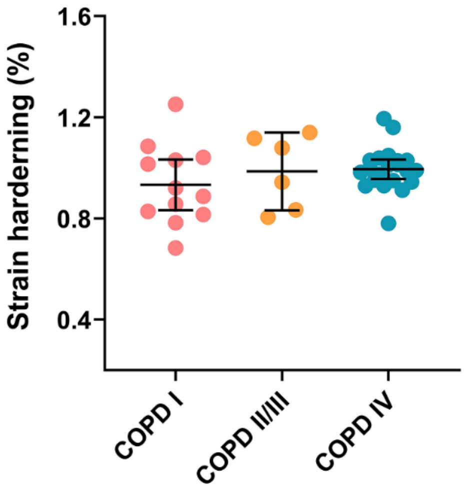


**Figure S5: Strain hardening of human COPD lung tissue.** Stress relaxation curves obtained from low load compression testing were used to evaluate strain hardening of COPD human lungs at different disease stages including stage I (n=4), stage II/III (n=2) and stage IV (n=8) and were plotted in triplicates per donor. Mean with 95% CI was plotted and One-way ANOVA with Dunnett's correction was used to evaluate differences between groups. A p-value <0.05 was considered significant. COPD: chronic obstructive pulmonary disease; dECM: decellularized extracellular matrix.
